# Random-walk based Probabilistic Microglia Morphology Reconstruction in 2D and 3D from Confocal Microscopy Images

**DOI:** 10.64898/2026.11.24.768701

**Authors:** Xinyi Chen, Ekram Hossain, Matthew N. McCall

**Affiliations:** University of Rochester Medical Center Rochester, NY, USA

## Abstract

Reconstructing individual microglia from confocal images is challenging when fine processes have weak fluorescence or neighboring cells form connected structures. Inspired by Grady’s random-walker framework, we propose a soma-guided probabilistic method for three-dimensional microglial reconstruction and cell-of-origin assignment. Signal-weighted random walks quantify voxel connectivity to each soma by the probability of reaching that soma before background or another soma. A path-length adjustment reduces the penalty on distal processes. Adjusted scores are compared across somata to assign voxels, subject to score and image-intensity thresh-olds. We assess process recovery and cell assignment separately using exact three-dimensional matching to manual tracings. In a P28 single-cell example, the reconstruction covered 83.98% of traced voxels within the analyzed region. For a connected cell pair in sample 12hrMD82, cover-age was 71.86% and 82.31%, with a pooled incorrect-assignment rate of 4.61% among detected traced voxels. Local analyses were also combined to produce an overview of 13 cell reconstructions in the same sample. We additionally provide a two-dimensional version that tolerates short signal gaps in maximum-intensity projections. These initial examples demonstrate the potential of soma-specific connectivity for microglial reconstruction.

## Introduction

Microglia are the resident immune cells of the central nervous system. They contribute to tissue surveillance, debris removal, and responses to injury and inflammation [2, 7]. Changes in the tissue environment can alter microglial morphology, including soma size, process length, and branching [2, 6]. These morphological features can be quantified to characterize microglial responses to injury [6].

Confocal microscopy is widely used to visualize microglial morphology in fluorescently labeled tissue. It collects two-dimensional images at different tissue depths to form a three-dimensional image stack. Resolution is typically higher within each image than along the depth (z) axis. Microglial morphology can be analyzed directly in the three-dimensional stack or in a two-dimensional maximum intensity projection.

However, accurately measuring these features often requires substantial image processing, including denoising, segmentation, and skeletonization, to isolate individual microglia from noisy images. Missing genuine processes can lead to underestimation of process length and branching. Including noise or false connections can lead to overestimation. Assigning a process to the wrong soma can also distort the morphological measurements of multiple neighboring cells.

Complete reconstruction of individual microglia remains challenging for two main reasons. First, microglial processes are long and thin, and their fluorescence signal often weakens with increasing distance from the soma. Distal processes may therefore have signal intensities that are hard to distinguish from background noise. Strict selection criteria can remove genuine processes, whereas permissive criteria can retain additional noise. Second, processes from neighboring microglia may touch in three dimensions or overlap in two-dimensional projections. Retaining depth information helps distinguish processes that overlap only in projection, reducing apparent connections between different cells. However, contacts between processes can still make cell assignment difficult in three-dimensional images. Detecting a process therefore does not necessarily identify the cell to which it belongs. Accurate reconstruction requires both the recovery of weak processes and their correct assignment to the soma of origin.

Existing methods often reconstruct each microglial cell as a tree extending from its soma. Megjhani et al. [5] used learned features of microglial processes to guide geodesic tree growth from multiple somata. Abdolhoseini et al. [1] identified seed points at multiple intensity thresholds and connected them into soma-rooted trees using geodesic paths. These methods provide a clear tree-like representation of cell morphology. However, only the selected connections are retained in the final tree, and alternative paths with similar image support are not directly represented. Weak processes may also be missed if suitable seed points are not detected or their connections are rejected during tree growth.

Another group of methods first converts the image into a binary cell mask and then extracts a skeleton from the segmented region. During this process, differences in fluorescence intensity are no longer directly represented: both strong and weak signals are treated as foreground. If processes from two neighboring microglia are connected in the binary mask, skeletonization represents them as a single connected network. Skeletonization alone does not determine the cellular origin of each branch, so an additional assignment rule is required to associate the branches with individual soma. Microglia Morphology Quantification Tool (MMQT) [4] addresses this problem by assigning each skeleton point to the soma with the shortest path along the skeleton. However, the nearest soma is not always the most strongly supported soma in the original image. A process may have a shorter skeletal path to soma A, while the route to soma B contains stronger and more continuous fluorescence signal. An assignment based mainly on skeletal distance may therefore assign the process to the wrong cell. Kimimaro [9] extracts skeletons from existing segmentations. When neighboring cells are merged in the segmentation, an additional step is needed to assign the resulting branches to individual somata. Some methods use morphological erosion to separate cells that are connected in a segmentation mask. MIC-MAC, developed by Salamanca et al. [8], uses iterative erosion to extract individual cells from connected segmented structures. Erosion can break narrow connections while leaving thicker regions connected. However, both genuine microglial processes and unwanted connections between cells can be thin. Thickness alone therefore cannot distinguish a genuine process from an unwanted connection.

To address these limitations, we developed a probabilistic reconstruction method based on the random-walker framework introduced by Grady [3]. The method first defines a candidate region using a permissive signal threshold. Within this region, a signal-weighted random walk measures the connectivity between each voxel and a target soma. The resulting hitting probability combines information from multiple possible paths through the image. Weak signals can therefore be evaluated using the signal intensities along their connections to a soma, as well as their local intensity.

We further extend this framework to distinguish processes belonging to neighboring microglia. For each voxel, the method compares its probabilistic connectivity to neighboring somata using random walks weighted by voxel signal intensities. Each voxel is assigned to a soma using the signal evidence along its connecting paths, rather than using skeleton path length alone. We introduce a path-length adjustment to prevent distant voxels with strong connecting signals from being excessively penalized by their longer paths. A voxel is assigned to the soma with the highest adjusted score only if this maximum is unique and both the score and normalized raw-image MIP intensity exceed their respective thresholds. Otherwise, the voxel remains unassigned.

The method combines evidence from multiple signal-supported paths to each soma, with a path-length adjustment to reduce the penalty on distal processes. We compare the adjusted scores for neighboring somata to assign processes within connected regions to individual cells. We also provide a two-dimensional version that can connect processes across short gaps in the observed signal. Process recovery and cell-of-origin assignment are evaluated separately using manual-trace coverage and incorrect-assignment rates. The reconstruction requires no additional training and allows different directional constraints on the random walk.

## Methods

### Method overview

Our method reconstructs individual microglia by comparing voxel connectivity to neighboring so-mata. The inputs are a preprocessed three-dimensional signal volume and identified soma targets. For each target, a signal-weighted random walk gives the probability of reaching that soma before reaching background or a competing soma. We adjust this hitting probability for geodesic path length to reduce the penalty on distal processes.

For a local region containing *n*_1_ target somata, each voxel *v* has a vector of connectivity scores:

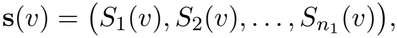

where *S*_*i*_(*v*) is the path-length-adjusted score for soma *i*.

For voxels outside the soma targets, assignment requires both the highest score and the normalized raw-image maximum-intensity projection (MIP) value to exceed their respective thresholds. A voxel meeting both thresholds is assigned to the soma with the unique highest score. All other voxels remain unassigned. Assigned voxels are combined with their soma targets to form cell-specific masks. Probability calculations and assignments are performed in three dimensions; the masks are projected for visualization.

### Soma-specific connectivity

To assess how strongly a voxel is connected to a target soma, we calculate the probability that a signal-weighted random walk starting at that voxel reaches the soma before reaching background or a competing soma. A higher probability indicates stronger connectivity under this model. This probability is defined by the active domain, transition probabilities, and soma and background boundary conditions.

#### Active domain

The active domain contains voxels considered as possible parts of the cells, while voxels outside it are treated as background. Candidate voxels satisfy *q*(*v*) ≥ *t*, where *t* is the active threshold, and are grouped into three-dimensional connected components. Two voxels are considered neighbors if they share a face, edge, or corner. This gives 26 possible neighbors within a 3 × 3 × 3 block after excluding the central voxel, allowing diagonally adjacent signal voxels to belong to the same component.

To identify components close to a soma, each soma mask is expanded by two rounds of three-dimensional dilation using the same neighborhood. A component is retained in full if any of its voxels overlaps an expanded mask. The retained components and the original soma targets form the active domain *A* ⊆ Ω, where Ω is the three-dimensional region of interest used for reconstruction. The expanded masks are used only for this contact test. Threshold selection is described under Implementation details.

#### Transition probabilities

To favor movement toward the target soma while allowing curved paths, we limit how far a single step can move away from the soma in the *xy* plane. Let *d*_*i*_(*v*) denote the Euclidean distance from voxel *v* to the center of target soma *i* in this plane. Among the 26 neighbors within the analysis region Ω, a neighbor *u* is eligible when

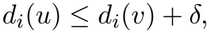

where *δ* is the allowed outward distance per step. This rule permits lateral movement and small outward steps rather than requiring every step to move closer to the soma. The eligible neighbors form the set *N*_*i*_(*v*).

Among the eligible neighbors, the walk is more likely to move to voxels with stronger signal. We define the transition weight and probability as

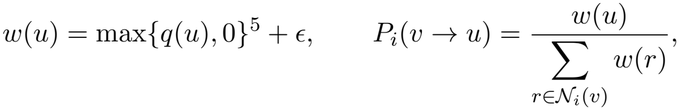

where *ϵ >* 0 prevents zero weights. The fifth-power transformation increases the contrast between strong and weak signal. The denominator includes all eligible neighbors, including those outside the active domain. These outside voxels have hitting probability zero, so a step into them counts as failure to reach the target soma.

#### Hitting probabilities and competing somata

For target soma *i*, the hitting probability *h*_*i*_(*v*) is the probability that a walk starting at voxel *v* reaches that soma before reaching another soma or leaving the active domain. Let *S*_*i*_ denote the set of voxels in the target soma mask for cell *i*. The boundary values are

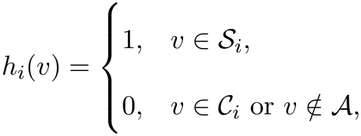

where *C*_*i*_ contains all voxels belonging to other detected somata within the analysis region. A walk stops when it reaches another soma or leaves the active domain. Either event counts as failure to reach the target soma.

A separate probability map is calculated for each soma to be reconstructed. In each map, the target soma is fixed at one and all other detected somata are fixed at zero, including those not being reconstructed in that analysis.

For an active voxel *v* outside the soma regions, the walk moves to a neighbor *u* with probability *P*_*i*_(*v → u*). From *u*, its probability of reaching the target soma is *h*_*i*_(*u*). Considering all eligible next steps gives

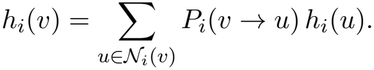

Thus, the hitting probability at *v* is a weighted average of its neighbors’ hitting probabilities. The probability map is obtained by solving these equations while keeping the soma and background boundary values fixed.

### Path-length adjustment and cell assignment

A low hitting probability can result from weak connecting signal or from a long path to the soma. Even along a signal-supported process, a longer walk has more opportunities to reach background or a competing soma. We therefore introduce a path-length adjustment to reduce the penalty on distal processes.

Let *ℓ*_*i*_(*v*) be the length of the shortest 26-neighbor path from voxel *v* to the target soma *S*_*i*_ within the active domain. Path length is the sum of the distances between consecutive voxels, measured in voxel coordinates. We use an effective length

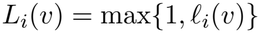

to avoid division by zero at the soma, and define the connectivity score

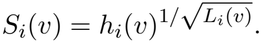

The square-root exponent provides a moderate adjustment. For the same hitting probability and a path length greater than one, it raises the score less than taking the *L*_*i*_(*v*)th root, 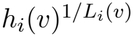. We use this partial adjustment to improve retention of distal signal while limiting the larger score increases produced by the latter transformation. Voxels with no path to the target or with zero hitting probability receive score zero. The adjusted score is used for cell assignment and is not interpreted as a probability of cell origin.

For a local region containing *n*_1_ target somata, the corrected scores at voxel *v* form a vector

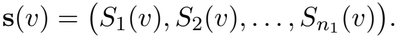

These scores are compared to determine cell identity.

For candidate voxels outside the soma targets, assignment requires a unique highest score above a threshold *τ*_*S*_. The normalized raw-image maximum-intensity projection (MIP), denoted by *I*(*x, y*), must also exceed an intensity threshold *τ*_*I*_. Thus, voxel *v* = (*x, y, z*) is assigned to cell *i* when

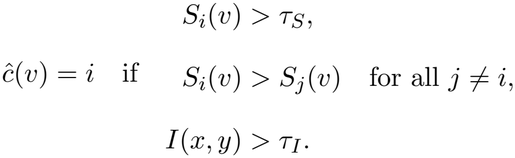

Voxels that fail either threshold or have tied maximum scores remain unassigned. Assigned voxels are combined with their corresponding soma targets to form cell-specific masks. The soma targets are always retained.

Hitting probabilities, path lengths, and cell assignments are calculated in three dimensions. For intensity filtering, all voxels at the same (*x, y*) position share the same raw-image MIP value, regardless of depth. The final cell masks are projected for visualization.

We also provide a two-dimensional version for maximum-intensity projections. For each soma, a signal-weighted random walk computes the probability of reaching the soma target within a local image region. The walk can cross short low-signal gaps, with a parameter *k* controlling how many consecutive low-signal steps are allowed before termination. We provide a brief demonstration of the two-dimensional method’s performance. Figure 1 illustrates this approach for six selected somata in sample `12hrMD82.`

**Figure 1:**
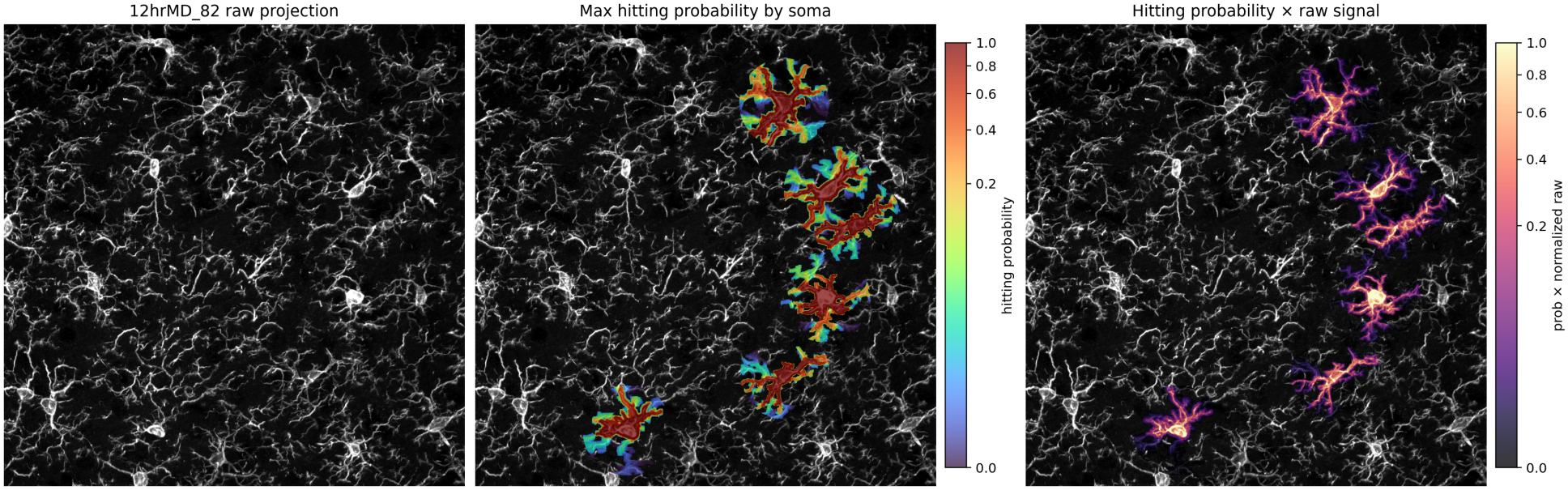
Two-dimensional results for six selected somata in sample `12hrMD82`. From left to right: the raw-image maximum-intensity projection, the maximum hitting probability across the selected soma targets, and this probability map multiplied by the normalized raw-image intensity.

However, when full image stacks are available, we recommend the three-dimensional method. Maximum-intensity projection removes depth information: processes that are separated along the *z*-axis can appear connected in the *xy* projection. A two-dimensional random walk may follow these apparent connections and include processes belonging to another cell. The three-dimensional method retains depth information and evaluates connectivity within the image stack.

Figure 1 shows the two-dimensional results for six selected somata in sample `12hrMD82.`

### Evaluation against manual tracing

We evaluate process recovery and cell assignment separately using manual tracings. Let *T*_*i*_be the set of unique traced voxels for cell *i* within the analyzed ROI, and *R*_*i*_ its final reconstruction mask. Define

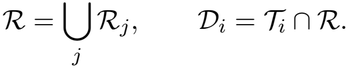

Thus, *D*_*i*_ contains traced voxels detected in any cell mask, regardless of whether their assigned identity is correct.

Coverage measures the fraction of traced voxels detected:

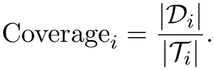

The denominator includes all traced voxels within the ROI, including those outside the active domain or left unassigned.

Among detected traced voxels, those outside the correct cell mask *R*_*i*_ are incorrectly assigned. The assignment-error rate is

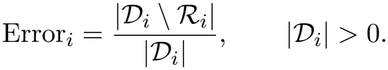

Pooled measures are calculated by summing the corresponding numerators and denominators across cells. All comparisons use exact three-dimensional voxel coordinates.

## Implementation for 3D

The reconstruction procedure was applied to single-cell, connected-pair, and multicell analyses. The overall workflow is shown in Figure 2. Input preparation, computational settings, and output processing are described below.

**Figure 2:**
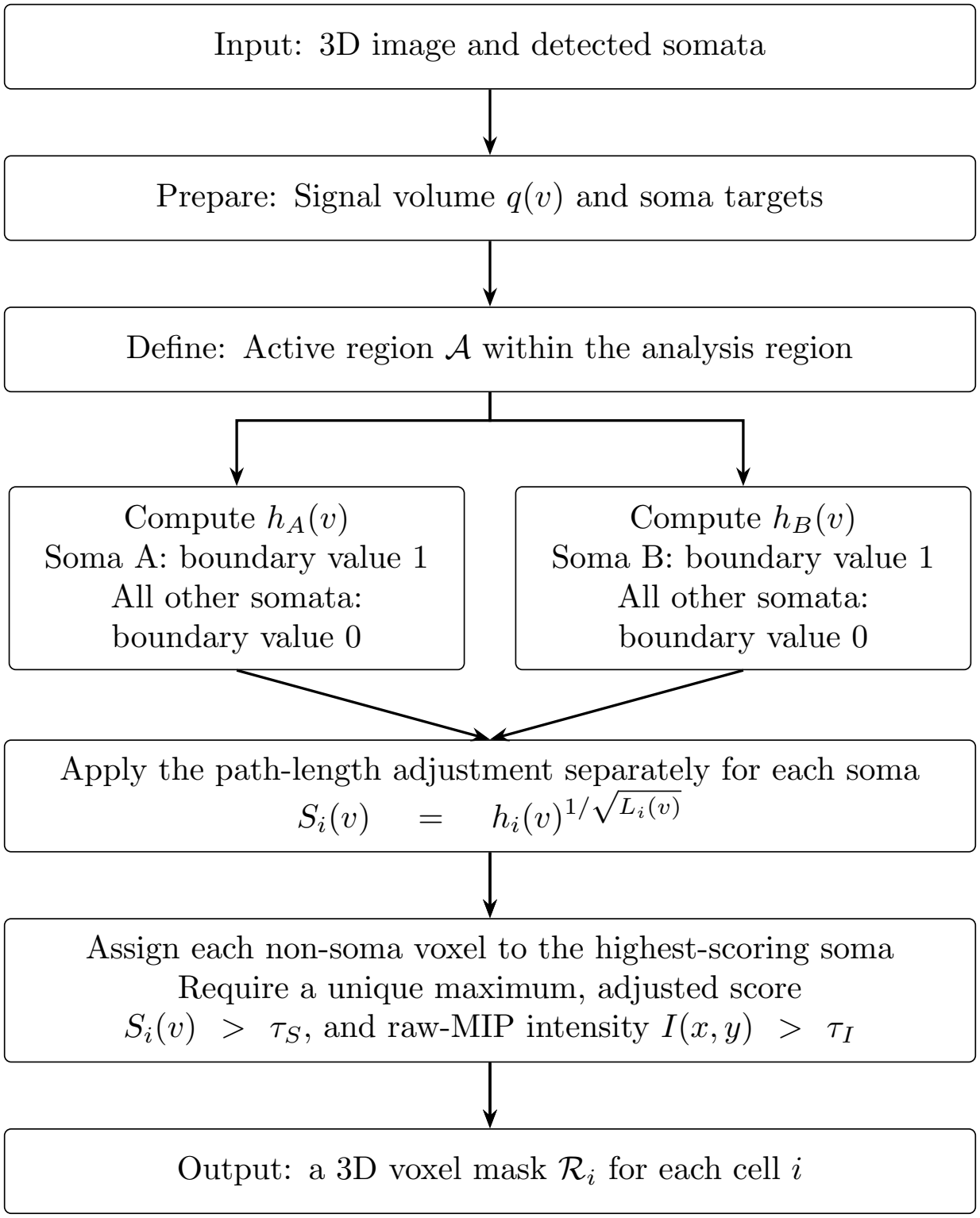
Cell Reconstruction workflow. Separate hitting-probability maps are computed for each target soma, illustrated here for somata A and B. In each calculation, the target soma has boundary value 1 and all other detected somata have boundary value 0. The probabilities are adjusted for geodesic path length and compared across target somata to determine voxel assignments. Voxels that fail either threshold or have tied maximum scores remain unassigned. Soma targets are retained throughout.

### Inputs and analysis regions

Following the distance-based weighting approach described in VonKaenel’s doctoral thesis [10], voxel intensities were multiplied by a spatial weight that decreased with distance from the detected somata. The resulting values were normalized to [0, 1].

For the manually traced examples, Cellpose labels provided the soma centers. Each soma target contained voxels at or above a specified signal percentile within an axial cylinder centered on the soma. The cylinder radius and percentile are listed in Table 1. For the multicell analysis of `12hrMD82`, existing three-dimensional soma masks were used directly.

**Table 1:**
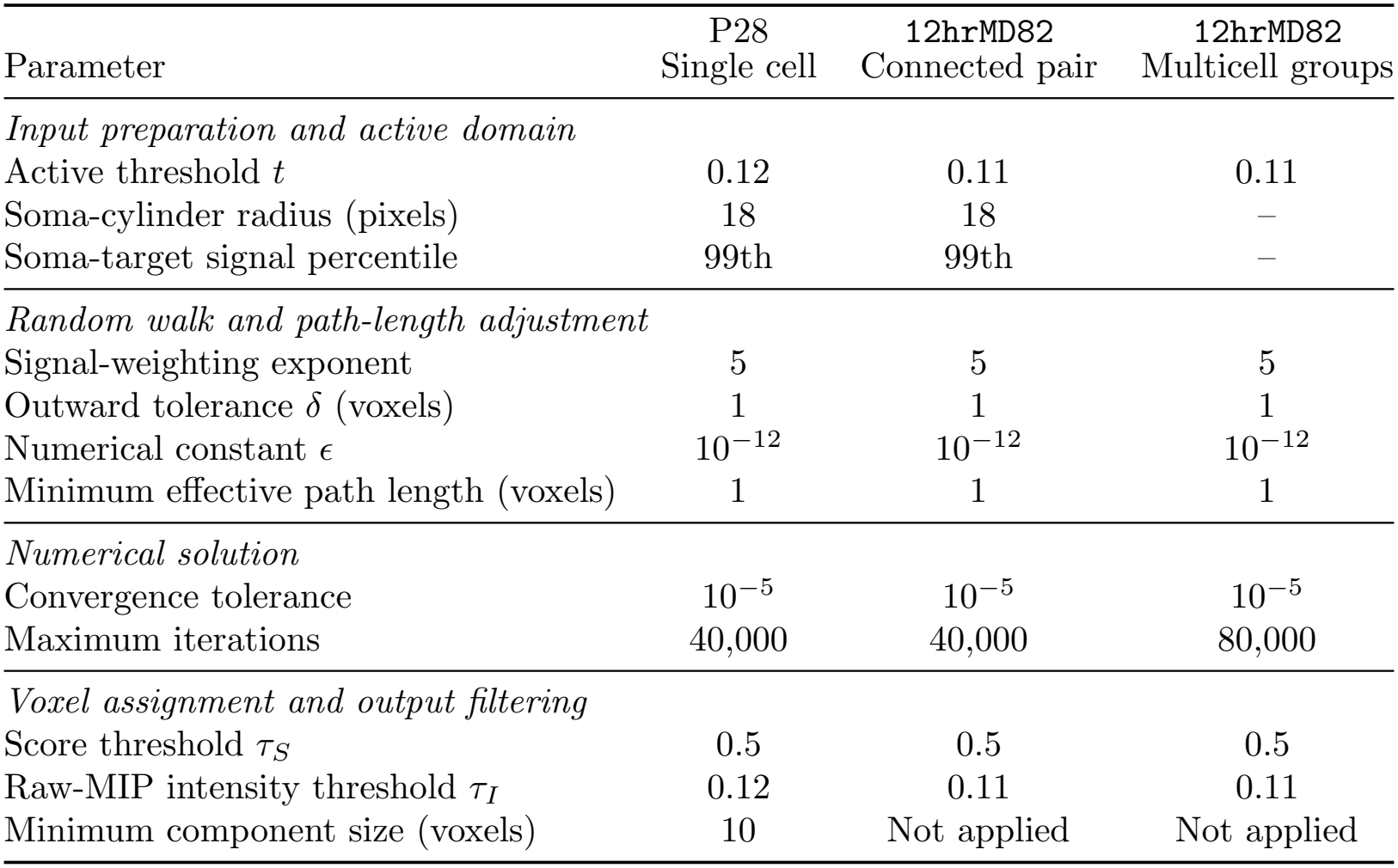
Settings used in the three-dimensional analyses, grouped by their role in the implementation. Dashes indicate settings that were not required because existing soma masks were used.

| Parameter | P28<br>Single cell | 12hrMD82<br>Connected pair | 12hrMD82<br>Multicell groups |
|---|---|---|---|
| *Input preparation and active domain* |  |  |  |
| Active threshold *t* | 0.12 | 0.11 | 0.11 |
| Soma-cylinder radius (pixels) | 18 | 18 | – |
| Soma-target signal percentile | 99th | 99th | – |
| *Random walk and path-length adjustment* |  |  |  |
| Signal-weighting exponent | 5 | 5 | 5 |
| Outward tolerance *δ* (voxels) | 1 | 1 | 1 |
| Numerical constant *ϵ* | 10<sup>−12</sup> | 10<sup>−12</sup> | 10<sup>−12</sup> |
| Minimum effective path length (voxels) | 1 | 1 | 1 |
| *Numerical solution* |  |  |  |
| Convergence tolerance | 10<sup>−5</sup> | 10<sup>−5</sup> | 10<sup>−5</sup> |
| Maximum iterations | 40,000 | 40,000 | 80,000 |
| *Voxel assignment and output filtering* |  |  |  |
| Score threshold *τ<sub>S</sub>* | 0.5 | 0.5 | 0.5 |
| Raw-MIP intensity threshold *τ<sub>I</sub>* | 0.12 | 0.11 | 0.11 |
| Minimum component size (voxels) | 10 | Not applied | Not applied |

The P28 cell was analyzed within a local region of interest (ROI), and the connected pair in `12hrMD82` was analyzed within a shared ROI. For the multicell analysis, neighboring somata were manually organized into local groups. Each group ROI retained all 41 axial slices and included a nominal 150-pixel margin in the imaging plane, clipped at the image boundaries. Results were returned to the original image coordinates without resolving overlaps between groups. Voxel competition was therefore restricted to each local analysis.

### Parameter settings and numerical computation of hitting probabilities

The reconstruction code was implemented in Python. Numba accelerated the iterative hitting-probability solver, and scikit-image provided the geodesic path-length calculation. The signal-weighting exponent, outward-step tolerance, and score threshold were held constant across analyses. The active-domain and raw-MIP intensity thresholds differed between samples. These thresholds act on the preprocessed signal *q*(*v*) and the normalized raw-image MIP *I*(*x, y*), respectively. Table 1 lists the settings used in each analysis.

The hitting-probability equations were solved by updating active non-soma voxels from the probability values at the previous iteration, while holding soma and background values fixed. Iteration stopped when the maximum absolute change between successive maps fell below the convergence tolerance or the iteration limit was reached.

Geodesic path lengths were computed using `MCP Geometric` from scikit-image, with all target-soma voxels supplied as starting points. Traversal costs were one within the active domain and infinite outside it. The calculation used all 26 neighbors, with geometric step lengths measured in voxel coordinates.

### Output masks and visualization

For the intensity gate, the raw-image maximum-intensity projection (MIP) was normalized separately within each ROI:

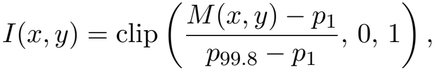

where *M* (*x, y*) is the maximum raw-image intensity along the axial direction, and *p*_1_ and *p*_99.8_ are the 1st and 99.8th percentiles of the MIP intensities within the ROI. The normalized values were used in the voxel-assignment rule defined in Methods.

After assignment, small components were removed from the P28 reconstruction using the size threshold in Table 1. All assigned components were retained in the connected-pair and multicell analyses. Soma targets were retained in all analyses.

For visualization, each cell’s corrected scores were projected along the axial direction by taking the maximum over its assigned voxels at each (*x, y*) position. The projections were overlaid on the raw-image MIP, with soma contours marking the targets. In combined views, colors indicated cell identity; in multicell displays, brightness indicated corrected score and colors were blended at projected overlaps. These overlaps can arise from structures at different depths. Visualization did not alter the three-dimensional assignments.

## Results

### Reconstruction of a manually traced microglial cell

We first evaluated the method on a manually traced microglial cell from a P28 sample `p28_330_HET_ 009`, using the manual tracing as the reference. This analysis examined whether the soma-specific hitting-probability framework could reconstruct the traced cell within a soma-centered region of interest (ROI), followed by assessing reconstruction quality using the coverage rate relative to the manual tracing ground truth.

Figure 3 shows the raw fluorescence image, the final reconstruction colored by the path-length-adjusted score 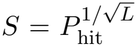 and the manual tracing within the same ROI. Here, *P*_hit_ is the soma-specific hitting probability and *L* is the path length to the target soma, as defined in the Methods. The reconstruction delineated the soma and multiple branching processes. In the projected score map, higher values of *S* were observed around the soma and along several major processes, whereas portions of the peripheral structure had lower scores.

**Figure 3:**
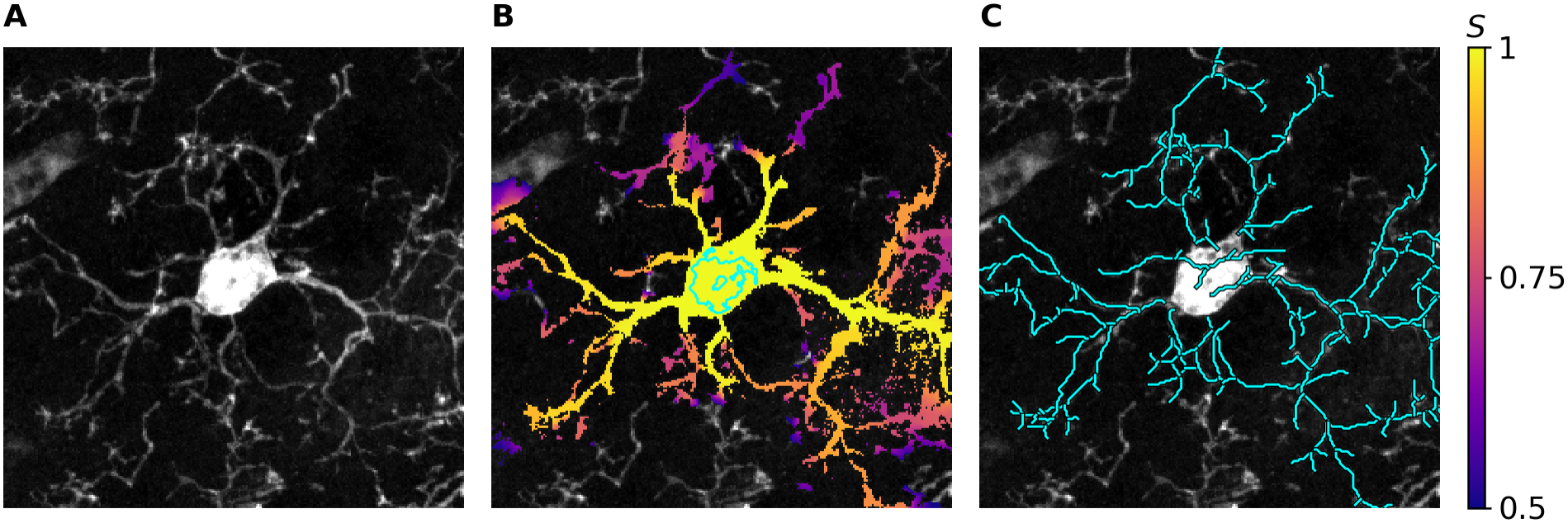
Reconstruction of a manually traced microglial cell (sample `p28_330_HET_009`). (A) Raw fluorescence maximum-intensity projection (MIP) within the analyzed ROI. (B) Reconstruction with adjusted connectivity score *S >* 0.5 and normalized raw-MIP intensity *I >* 0.12. Brighter colors indicate higher scores; the cyan contour marks the soma target. (C) Manual tracing shown in cyan over the same raw-image MIP.

To quantify agreement with the ground-truth reference, we evaluated whether each manually traced voxel within the ROI was included in the final reconstruction mask. The reconstruction covered 2,092 of 2,491 unique traced voxels, yielding an exact three-dimensional coverage of 83.98%. We additionally evaluated coverage after projection onto the imaging plane. Of the 2,399 unique traced pixels in this projection, 2,099 were covered by the reconstructed mask, corresponding to a projected coverage of 87.49%. Coverage was higher after projection because depth information was ignored. We therefore used exact 3D coverage to assess reconstruction quality. The final reconstruction retained most of the manually traced structure within the ROI. However, some noise on the right side of the ROI remained after filtering.

### Effect of path-length correction on reconstruction

We examined the effect of path-length correction in the same P28 ROI by comparing three scores: the uncorrected hitting probability *P*_hit_, the length-normalized score 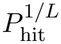, and the square-root-adjusted score 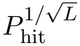. All three were evaluated using the same score threshold of 0.5, normalized raw-MIP intensity threshold of 0.12.

As shown in Figure 4, fewer peripheral processes were retained without path-length correction. A voxel far from the soma can have a low hitting probability even when it is bright and connected to the soma by a visible process. A random walk starting farther from the soma has more opportunities to enter the background before reaching it. This introduces a distance-dependent penalty that can suppress distal processes too strongly.

**Figure 4:**
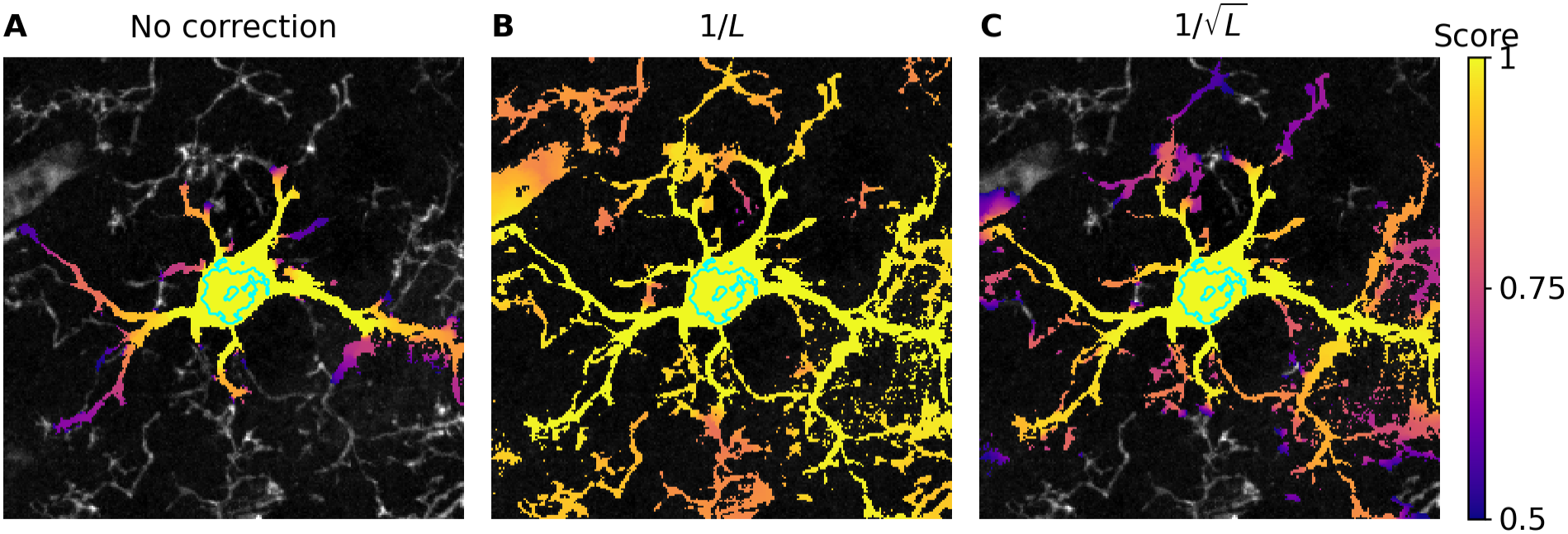
Effect of path-length correction on reconstruction (sample `p28 330 HET 009`). (A) Uncorrected hitting probability, *Q* = *P*_hit_. (B) Length-normalized score, 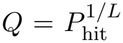. (C) Square-root-adjusted score, 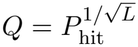. All three reconstructions used *Q >* 0.5 and normalized raw-MIP intensity *I >* 0.12. Components smaller than 10 voxels were removed, while components containing soma targets were retained. Colors show the maximum score among selected voxels along the axial direction, overlaid on the same raw-image MIP. Brighter colors indicate higher scores. Cyan contours mark the projected soma target.

We therefore introduced path-length correction to relax this distance-dependent penalty. We considered two forms: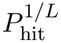 and 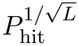, where *L* is the geodesic path length to the soma. We can illustrate the difference between these corrections using a simplified exponential-decay model, *P*_hit_ *≈ e*^*−aL*^, where *a* > 0 is the decay rate. Taking the *L*th root gives 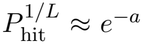, removing the dependence on path length in this model. In Figure 4, this correction raised scores across a broad portion of the ROI, suggesting that it relaxed the distance penalty too strongly.

The square-root correction provides a more moderate relaxation. Under the same model, 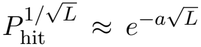, the scores still decrease with path length, but more slowly. In this example, it retained more peripheral structure than the uncorrected map while avoiding the extensive expansion produced by the 1*/L* correction. We therefore used the square-root correction as a fixed partial relaxation of the distance penalty, without fine-tuning the exponent against the manual tracing.

### Separation and cell-of-origin assignment of connected microglial cells

We next evaluated the method on two connected microglial cells in the contralateral primary visual cortex (V1) following 12 hours of monocular deprivation (sample `12hrMD82`). Manual tracings of both cells served as the ground-truth references. This analysis addressed a different challenge from single-cell reconstruction: whether processes within a shared connected region could be assigned to the correct soma.

Both cells were analyzed within a shared ROI (Figure 5A). We computed a separate hitting-probability map for each soma. When computing the map for Cell A, we set the probability to 1 at the soma of Cell A and 0 at the soma of Cell B, and reversed these values when computing the map for Cell B. After path-length correction, voxels passing the score and raw-image intensity thresholds were assigned to the cell with the higher corrected score. The resulting cell-specific masks are shown separately in Figure 5B,C and together in Figure 5D. These masks did not overlap in three dimensions, although processes at different depths could overlap in the two-dimensional projection.

**Figure 5:**
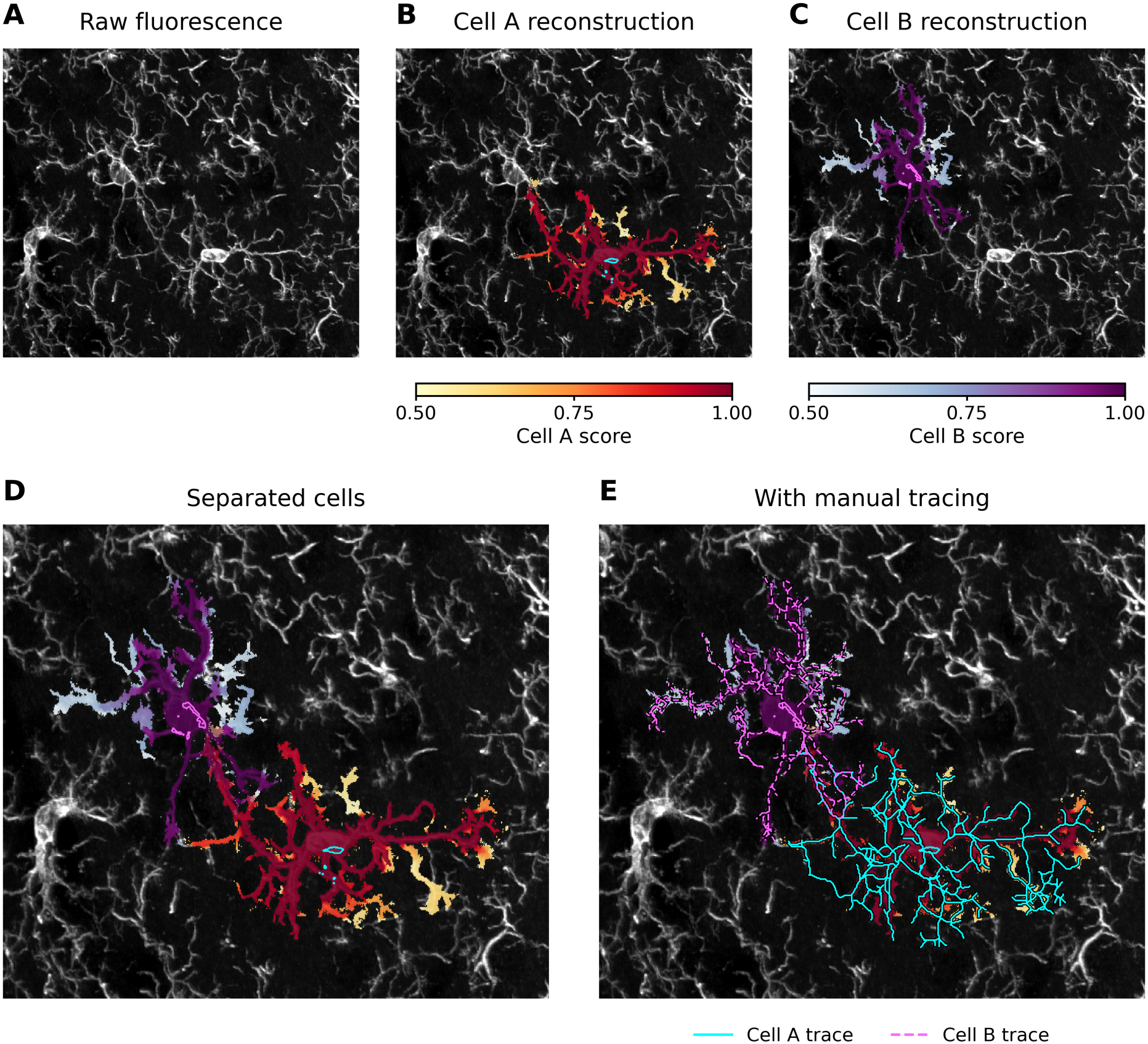
Reconstruction and cell-of-origin assignment of two connected microglial cells in sample `12hrMD82`. (A) Normalized raw-image maximum-intensity projection (MIP) within the shared ROI. (B,C) Reconstructions of Cell A and Cell B, respectively. Colors indicate the maximum assigned path-length-adjusted score *S* along the axial direction. (D) Combined projection of the two reconstructions; colors are averaged where their projections overlap. (E) The same reconstructions with manual tracings overlaid: cyan solid lines for Cell A and magenta dashed lines for Cell B. Cyan and magenta contours mark the corresponding soma targets. Assignment required a unique highest score with *S >* 0.5 and normalized raw-MIP intensity *I >* 0.11. Soma targets were retained independently of these thresholds, and no small-component filtering was applied. Coverage and incorrect-assignment rates were evaluated using exact three-dimensional voxel matching to the manual tracings within the ROI.

We compared the reconstructions with the manual tracings (Figure 5E) and quantified coverage and cell assignment using exact three-dimensional matching within the ROI. For Cell A, 2,142 of 2,981 traced voxels were detected (71.86%). For Cell B, 1,935 of 2,351 traced voxels were detected (82.31%). Across both cells, 188 of the 4,077 detected traced voxels were assigned to the incorrect cell, giving a pooled misclassification rate of 4.61%. Thus, most detected reference locations were assigned correctly, while incomplete coverage remained the larger source of disagreement with the manual tracings.

### Multicell reconstruction in sample 12hrMD82

We extended the analysis beyond the manually traced pair to 11 additional MGtoolbox-detected somata in sample `12hrMD82`. Neighboring somata were manually organized into five local groups.

This grouping assumed that distant groups without evident signal-supported connections had limited influence on one another’s reconstruction. Local ROIs reduced the size of each probability calculation and avoided repeatedly processing distant image regions.

Groups 1 and 5 illustrate this approach. Group 1 contained somata 1, 2, and 4 in the upper portion of the image (Figure 6A), whereas Group 5 contained somata 3, 11, and 7 on the right side (Figure 6A). Each ROI enclosed its group with a nominal margin of 150 pixels in the imaging plane and retained all 41 axial slices. After truncation at the image boundaries, the ROI dimensions were 576 × 395 × 41 voxels for Group 1 and 278 × 473 × 41 voxels for Group 5, in *X, Y, Z* order. Both groups used an MG active threshold of 0.11, a corrected-score threshold of 0.5, and a normalized raw-MIP intensity threshold of 0.11.

**Figure 6:**
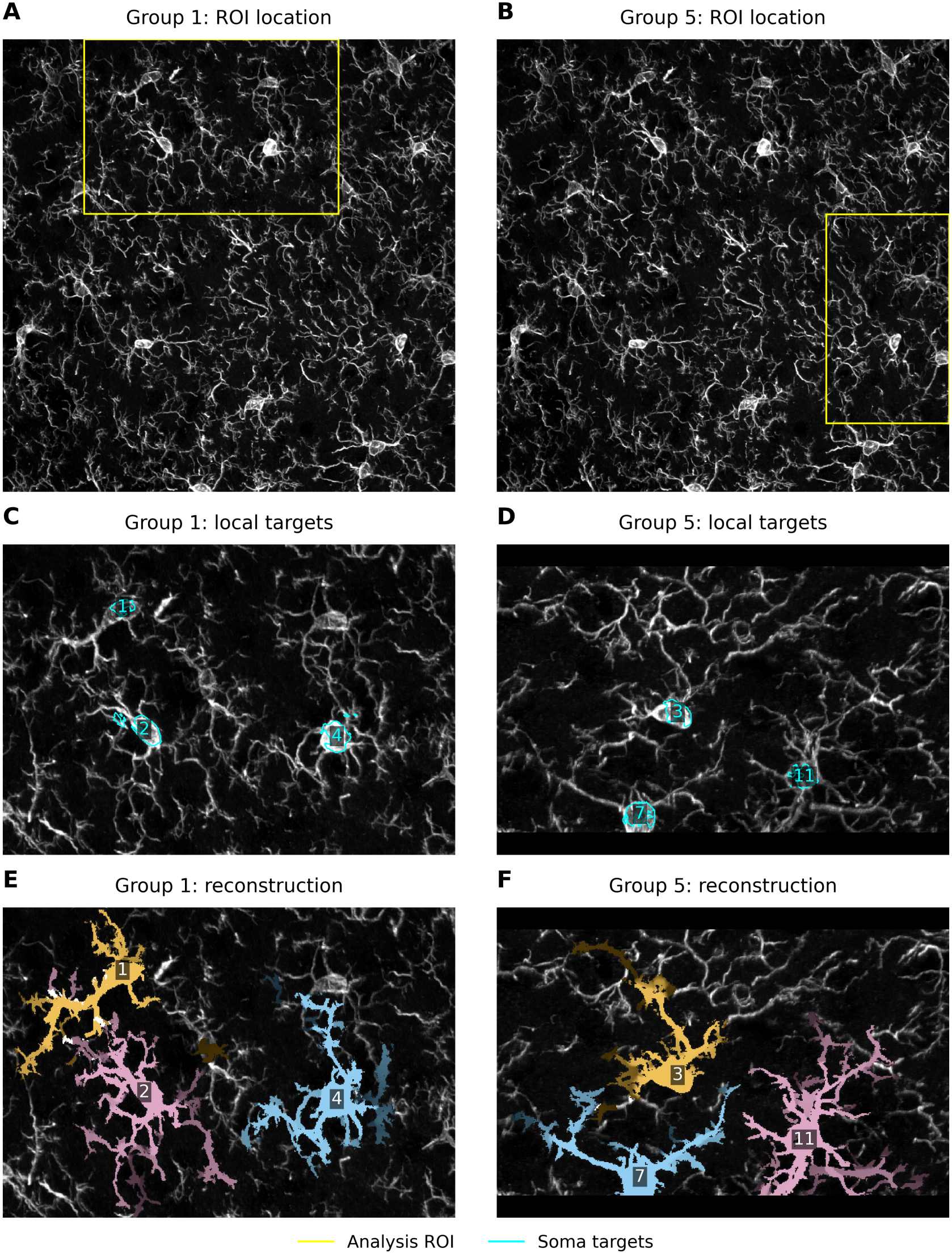
Local reconstruction of Groups 1 and 5 in sample `12hrMD82`. (A,B) Full-image MIPs with the corresponding analysis ROIs outlined in yellow. (C,D) Local MIPs with soma targets outlined and numbered in cyan: somata 1, 2, and 4 for Group 1, and somata 3, 11, and 7 for Group 5. (E,F) Corresponding reconstructions. Different hues identify individual cells, and brighter shades indicate higher maximum assigned adjusted scores *S* along the axial direction, over the displayed range of 0.5–1. White marks overlaps between cell projections. Panels D and F were rotated 90° clockwise and padded to match the dimensions of C and E without stretching or cropping.

The reconstructions retained branching structures around the individual somata and distinguished their assignments within each shared ROI (Figures 6B and 6B). Different color families indicate individual cells, while brighter shades represent higher corrected scores. All six soma-specific robability calculations converged with a maximum successive-update difference below 10^−5^.

Reconstructions from the five local groups and the previously analyzed cell pair were mapped to their original image coordinates, yielding an overview of 13 reconstructed cells (Figure 7). Voxel assignments from each local analysis were retained, and overlaps between separate analyses were not resolved.

**Figure 7:**
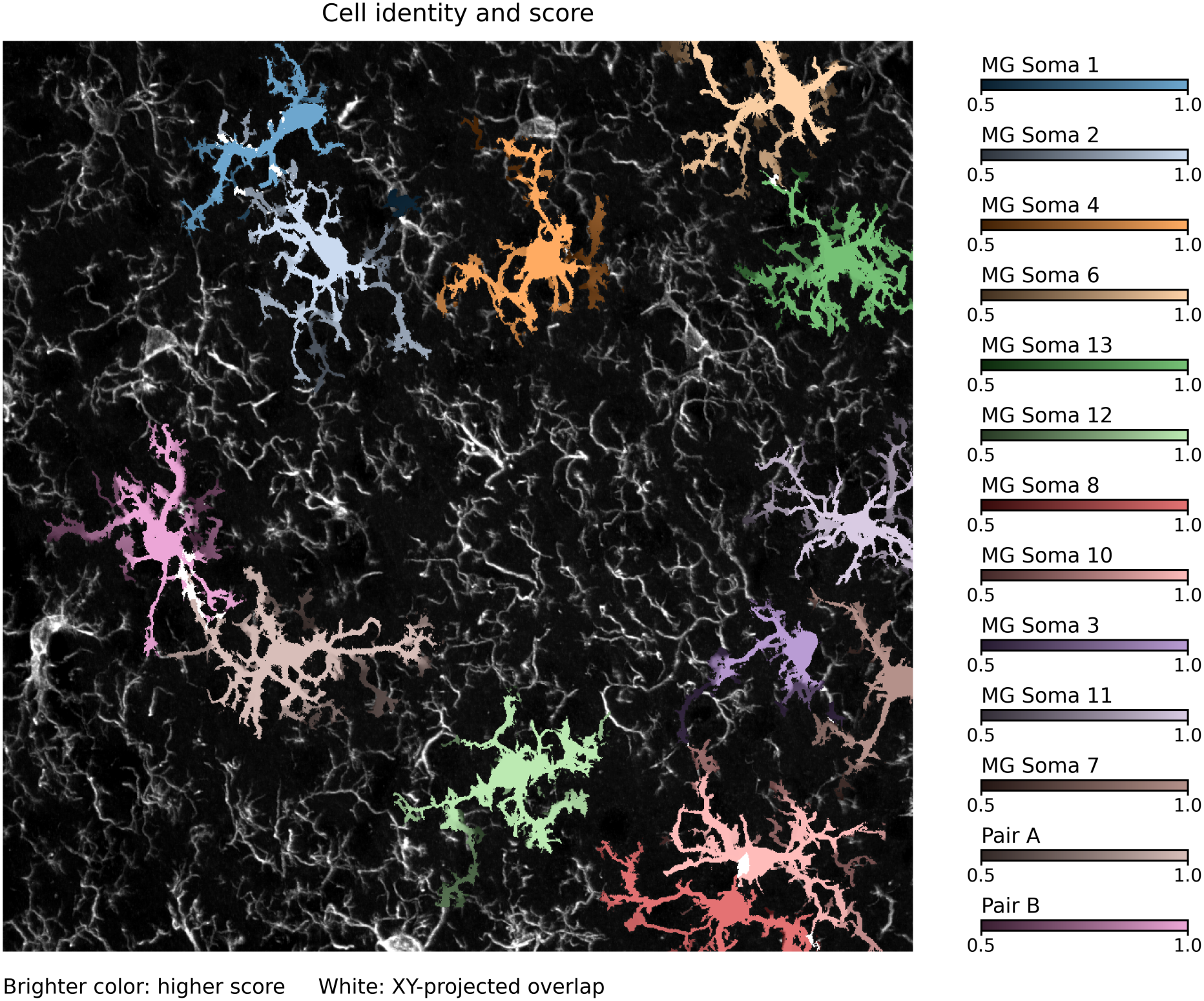
Full-image overview of 13 cell reconstructions in sample `12hrMD82`, combining five local groups with the previously analyzed cell pair. Different hues identify individual cells, and brighter shades indicate higher maximum assigned adjusted scores along the axial direction. White marks pixels containing multiple cell projections. Local reconstructions were placed in their original image coordinates, retaining their original cell assignments and overlaps between separate analyses.

## Discussion and limitations

We developed a method for reconstructing individual microglia and assigning their processes to somata in three-dimensional confocal images. Signal-weighted random walks measure voxel-to-soma connectivity by combining evidence from multiple paths through the image. This allows cell assignment to use the fluorescence signal along the connecting paths. A path-length adjustment reduces the penalty on distal processes, and the adjusted scores are compared across somata to determine voxel assignments.

In the P28 sample, the final reconstruction covered 83.98% of the manually traced voxels within the analyzed ROI. For the connected cell pair in `12hrMD82`, coverage was 71.86% and 82.31%, with a pooled misclassification rate of 4.61% among detected traced voxels. Together, these results suggest that signal-weighted connectivity can support both recovery of cell structure and assignment of processes to individual somata.

Path-length correction helped retain peripheral structures that received low uncorrected hitting probabilities. Even when a distal process is bright and connected to the soma through a signal-supported path, a longer random walk has more opportunities to enter the background before reaching the soma. The resulting distance penalty can therefore suppress distal processes. In the P28 example, square-root correction retained more peripheral structure than the uncorrected map, while producing less extensive selection than normalization by *L*. This provides a practical partial relaxation of the distance penalty, although it can also increase scores in regions that do not belong to the target cell.

The connected-cell analysis showed that most detected tracing voxels were assigned to the correct cell. However, this assignment performance applies only to detected locations and does not account for processes that were missed entirely. Some fine processes and complex local structures identified by manual tracing were not fully recovered. Assigning a probability to each voxel does not guarantee that the thresholded reconstruction will preserve every detail of the traced structure. Further improvements should therefore aim to recover missing processes while limiting additional regions and maintaining accurate cell assignment.

The current validation is also limited to one manually traced cell in P28 and a connected pair in `12hrMD82`. These examples address single-cell recovery and connected-cell assignment, but do not establish performance across a broad range of imaging conditions or cell morphologies. In addition, tracing coverage measures how much reference structure is recovered; it does not determine whether all additional selected regions are correct. Broader validation with additional manually annotated cells is needed to assess the generalizability and specificity of the method.

The three-dimensional implementation requires iterative computation until the hitting probabilities converge, which can be computationally demanding for large images. The two-dimensional version described in Methods provides an option when only maximum-intensity projections are available. However, projection removes depth information. Processes separated along the *z*-axis may appear connected in the *xy* projection, allowing the random walk to follow an apparent connection between different cells. We therefore recommend the three-dimensional method whenever full image stacks are available.

## Conclusion

We developed a soma-guided probabilistic method for three-dimensional microglial reconstruction and cell-of-origin assignment. Signal-weighted random walks measure connectivity to individual somata, and a path-length adjustment reduces the penalty on distal processes. In the evaluated three-dimensional examples, the method recovered much of the manually traced structure and correctly assigned most detected tracing voxels to their soma of origin. We also provide a two-dimensional version for maximum-intensity projections, although the three-dimensional method is preferred when full image stacks are available. Further work is needed to improve recovery of fine processes, limit non-target regions, and reduce computational cost.

## Data and Code Availability

The microscopy images, manual tracings, and code used for microglial reconstruction, evaluation, and figure generation are publicly available at https://github.com/mccall-group/Random_Walk_Microglia.

## Notes

https://github.com/mccall-group/Random_Walk_Microglia

### Competing Interest Statement

The authors have declared no competing interest.

### Funding

- National Institute of Neurological Disorders and Stroke: R01NS136626

